# A switch from TE-like heterochromatin to euchromatin underlies activation of protein storage genes in maize endosperm

**DOI:** 10.1101/2025.09.26.678861

**Authors:** Yirui Sun, Yibing Zeng, Dong won Kim, R. Kelly Dawe, Jonathan I. Gent

## Abstract

Unlike the stable transcription of plant genes with CG gene body methylation alone, genes with methylation in TE-like CHG and CG contexts are poorly transcribed and often missanotated. In contrast, here we describe a set of TE-like methylated genes that are maternally demethylated and expressed toward the extreme high end of the spectrum in endosperm. They are enriched for short, secreted proteins, and about a third encode zeins, the major seed storage proteins of maize. Consistent with their dynamic expression in endosperm, they acquire either activating or repressive gene regulatory modifications when demethylated but are heterochromatic when methylated in other tissues. The majority are expressed both maternally and paternally, but a subset whose methylation/demethylation extends upstream of gene bodies into promoters are strongly imprinted. These and other data indicate that TE-like methylation and associated heterochromatin can be a signature of broad silencing but exceptionally high and specific gene expression in either pollen or endosperm.

## INTRODUCTION

While in many plants endosperm is an ephemeral tissue with little nutritional value in the mature seed, in cereal crops it provides a rich source of starch and a significant source of protein. That, combined with its ease of storage and transport, has made it the major source of the world’s calories (Sabelli and Larkins 2009). Placing it into the plant life cycle poses a problem: It is neither sporophyte nor gametophyte. Like the embryo, it is produced by fusion of two gametes. But unlike the embryo, it is a terminal side branch from the alternation-of-generations life cycle (Fig. 1A). Both male and female gametophytes carry two gametes, the male sperm cells, and the female egg cell and central cell. In most angiosperms the central cell has two haploid nuclei, and fertilization gives rise to a triploid endosperm (Fig. 1B) (Schmid et al. 2015). Its initial development is coenocytic, with rapid and synchronized nuclear replication without cell division producing up to 512 nuclei within the space defined by the central cell’s large cytoplasm. It is critical to embryo development and germination and in many species, including maize, to early seedling growth. Central to this is its role in transferring nutrients from maternal cells to the developing embryo and storing them for the young seedling during germination.

**Fig. 1:**
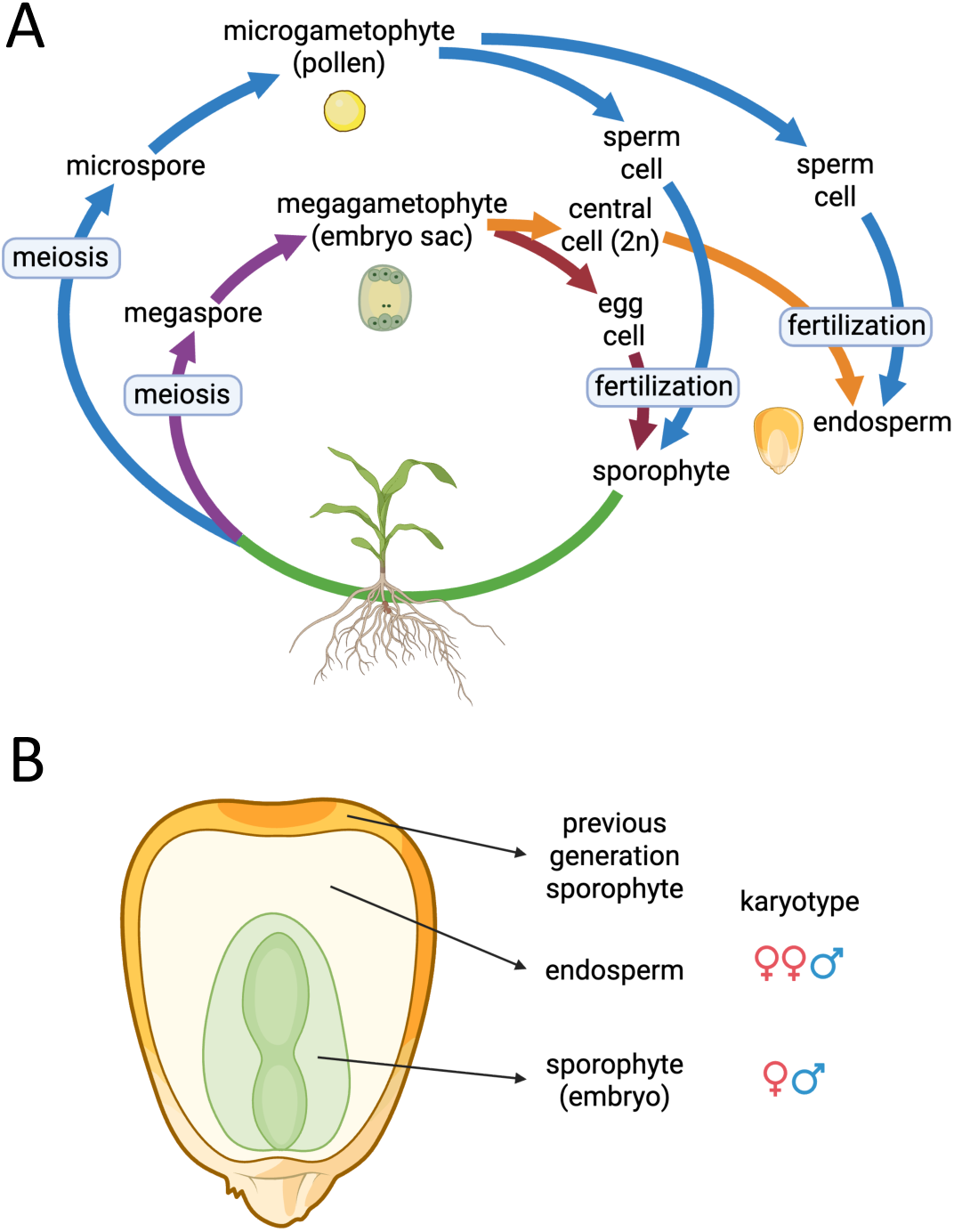
Endosperm, sporophyte, and gametophyte in the context of an angiosperm life cycle. **a** Endosperm is a terminal offshoot from the angiosperm lifecycle formed by fusion of sperm with a binucleate central cell (female gamete). **b** The three major parts of a kernel of maize. Endosperm is triploid because it inherits two copies of the maternal genome and one of the paternal genome. The previous generation sporophyte is called the pericarp in maize (not to scale).

In addition to the expected two-to-one ratio of maternal-to-paternal gene dosage, maternally imprinted genes can have much higher ratios of maternal-to-paternal dosage. In effect, this causes imprinted genes to maintain a partially haploid-like expression state after fertilization. The phenomenon is closely connected to repression of transposons/transposable elements (TEs). Not only are core mechanisms of TE silencing shared with mechanisms of imprinting, many TEs are themselves imprinted (Del Toro-De León et al. 2024; Rodrigues et al. 2021; Li et al. 2023; Pignatta et al. 2014; Anderson et al. 2021). In fact, the first example of an imprinted gene, the maize gene *r1*, was later discovered to have a CACTA DNA transposon as its promoter (Kermicle 1970; Walker 1998). In the best-studied form of imprinting in plants, DNA demethylation of maternally inherited cis-regulatory elements allows binding of activators and preferential maternal expression. Other forms of imprinting are also common (Batista and Köhler 2020). Imprinting by demethylation involves both CG and CHG methylation together (H = A,T, or C). Depending on the species and on other details of local DNA sequence context and chromatin context, it can also involve CHH methylation (Zemach et al. 2013; Gouil and Baulcombe 2016; Stroud et al. 2014). In maize, CHH methylation occurs with CHG and CG methylation at sites of RNA-directed DNA methylation (RdDM) at euchromatin/heterochromatin boundaries and has little connection to imprinting (Fu et al. 2018; Gent et al. 2022).

CG and CHG together, and in some cases CHH methylation, are a central feature of heterochromatin, where they function with histone modifications, histone variants, and other factors to repress transcription. DNA methylation does so primarily in the context of genome defense (e.g., TE silencing), and its repression is characteristically stable throughout development. This stability of DNA methylation throughout sporophyte development has been well-documented in maize (Crisp et al. 2020; Oka et al. 2017; Hufford et al. 2021). In some cases, however, targeted DNA demethylation can facilitate activation of gene expression, as occurs in some imprinted genes with methylated promoters. The enzymes responsible are bifunctional DNA glycosylases/lyases (DNGs for short) (Du et al. 2023). “Glycosylase” refers to the enzymes’ ability to cleave the sugar-base bond to create an abasic site, and “lyase” to their ability to cleave the sugar-phosphate backbone. The end result is base excision repair with an unmethylated cytosine. Mutant screens in Arabidopsis identified the DNG gene *DEMETER* as essential for imprinting of many genes, and mutants of a rice homolog showed a similar imprinting phenotype (Ono et al. 2012; Choi et al. 2002). A maize imprinting mutant identified soon after the discovery of *r1* imprinting called *maternal derepression of r1* (*mdr1*) also proved to encode a DNG (Kermicle 1978, 1995; Gent et al. 2022; Xu et al. 2022).

DNGs are critical to endosperm development as evidenced by early lethal mutant phenotypes, likely arising from failure to properly express master regulators of endosperm development (Ando et al. 2023; Haun et al. 2007; Tonosaki et al. 2021; Gehring et al. 2006; Hermon et al. 2007). Demethylation initiates prior to fertilization in Arabidopsis and rice, which could explain maternal specific demethylation (Park et al. 2016). DNG activity, however, is not limited to the central cell. In sporophytic tissues they function in methylation homeostasis and remove ectopic methylation from cis regulatory regions (Williams et al. 2022; Halter et al. 2021; Xu et al. 2025). In Arabidopsis, DNGs can target paternal alleles of some loci in endosperm (Hemenway and Gehring 2025) and activate gene expression in pollen (Borg et al. 2021; Khouider et al. 2021). DNG activity might also continue to operate on the maternal genome after merging of the two genomes. In sporophytic tissues DNGs target regions marked by histone H3 lysine 4 trimethylation (H3K4me3) to prevent ectopic methylation (Wang et al. 2025b). Possibly related to this, DNGs can interact with the histone variant H2A.Z (Nie et al. 2019). The factors that direct DNGs to demethylate specific imprinted loci in endosperm have not been identified, but as in the sporophyte, there is an association with transposons near genes (Tang et al. 2016; Frost et al. 2018; Zhang et al. 2019; Hemenway and Gehring 2025).

In contrast to methylation in promoters, methylation in gene bodies does not have a clear relationship to transcriptional repression. In maize as in other flowering plants, its effects depend on both local nucleotide contexts of the cytosines and larger contexts of genes and transposons (Zeng et al. 2023). CG and CHG methylation have little if any repressive effect on the genes that host them when they mark TEs in introns. An extreme example is the TR-1 kinesin gene, which contains over a hundred Kb of TEs in its introns, yet is actively transcribed (Swentowsky et al. 2020). CG and CHG methylation in coding exons, however, is strongly associated with poorly expressed, non-conserved, and misannotated genes (Zeng et al. 2023). Because of its similarity with methylation in TEs, this form of gene body methylation is referred to as TE-like methylation, abbreviated as teM (Kawakatsu et al. 2016). Most accurately annotated and functional maize genes lack CHG methylation (and lack CHH methylation) in their coding DNA but can have a spectrum of CG methylation levels correlated with constitutive and stable expression (Zeng et al. 2023). Genes on the higher end of the CG methylation spectrum, are called gene body methylated, abbreviated as gbM (Muyle et al. 2022).

While teM genes are largely silent or poorly expressed in sporophyte cells, a possible exception was observed in pollen early on, where some teM genes have robust pollen expression (Schmitz et al. 2013). (At the time, the term “RdDM targets” was used because the term “teM” had not been adopted yet, nor the subtleties of RdDM and non-RdDM CHH methylation understood). In a maize double mutant of two DNG homologs *mdr1* (a.k.a. *dng101*) and *dng102*, a set of ∼50 identified target genes exhibit ∼100-fold reduced expression in pollen relative to wild-type (Zeng et al. 2024). These genes, which are not the complete set of DNG targets in pollen, account for over 10% of the pollen transcriptome because of their unusually high expression levels in pollen.The majority are teM genes and have near-zero expression in sporophytic tissues. They are also unusual in their lack of introns, with most having either none or one. The majority have predicted functions in cell walls. Their high expression and pollen-specificity in pollen led us to ask whether there are similar teM genes with high expression in endosperm, where DNGs are already known to regulate imprinted gene expression. To answer this question, we used both gene expression patterns and TE-like methylation to define a list of 67 candidate genes. They encompass a wide variety of genes expressed at different stages of endosperm development, from secreted antifungal proteins and clavata homologs to transcription factors. Most strikingly, however, was the 23 zeins, the major seed storage prolamin genes of maize. These findings illustrate two broad types of DNG target genes in endosperm, classical imprinted genes with flanking methylation and moderate expression and tissue specificity, and the TE-like methylated genes with both flanking methylation and internal methylation across their gene bodies and unusually high expression and specificity. Consistent with their constitutive silencing throughout the life cycle of the plant, the TE-like methylated genes also had TE-like chromatin profiles in sporophyte tissues but gene-like profiles in endosperm. A subset of the TE-like methylated genes had imprinted expression, which could be attributed to methylation around transcription start sites rather than in gene bodies.

## RESULTS

### Identification of endosperm genes with TE-like methylation

Using similar methods as in previous studies, we defined teM genes as having both mCG and mCHG levels each equal or greater to 40% in coding DNA in leaf as a representative sporophyte tissue (Zeng et al. 2023, 2024). Since selecting for TE-like methylated genes also selects for mis-annotated genes, we only included core genes in this analysis, the ones that are annotated in syntenic positions in the B73 genome and all 25 other NAM founder genomes (Hufford et al. 2021). Given the temporal dynamics of gene expression through endosperm development, we defined each gene’s expression by its maximum TPM value over a time course of endosperm development (Chen et al. 2014). We then compared this value to its maximum expression in the nine non-endosperm tissues assayed in the NAM founder genome project. Using a minimum threshold of 20 TPM in endosperm and fivefold more expression in endosperm than any of the nine other tissues, we identified 67 genes that we refer to as endosperm teM genes.

### Reduced methylation of teM genes in endosperm

If endosperm teM genes are DNG targets, they should be methylated in sporophyte but demethylated in endosperm (at least for the two maternal genome copies). Comparisons of methylation profiles in endosperm and multiple sporophyte tissues showed a decrease in methylation in their gene bodies and in both flanks in endosperm (Fig 2A-C). For these comparisons, we used previously published EM-seq data from embryos and endosperm collected 15 days after pollination (15-DAP) from W22 inbred stocks (Gent et al. 2022), which required finding homologous gene annotations between W22 and B73 genomes. 42 of 67 endosperm teM genes had syntenic annotations in both genomes. The differences between sporophyte and endosperm were clearest in CG methylation because endosperm has a global reduction in CHG methylation that is not limited to DNG targets (Fig. 2C). CHH methylation was low in and near endosperm teM genes, similar to other teM genes (Zeng et al. 2023). We also compared endosperm with EM-seq data from W22 pollen (Zeng et al. 2024) and found no evidence for demethylation of endosperm teM genes in pollen (Fig 2A-C).

**Fig. 2:**
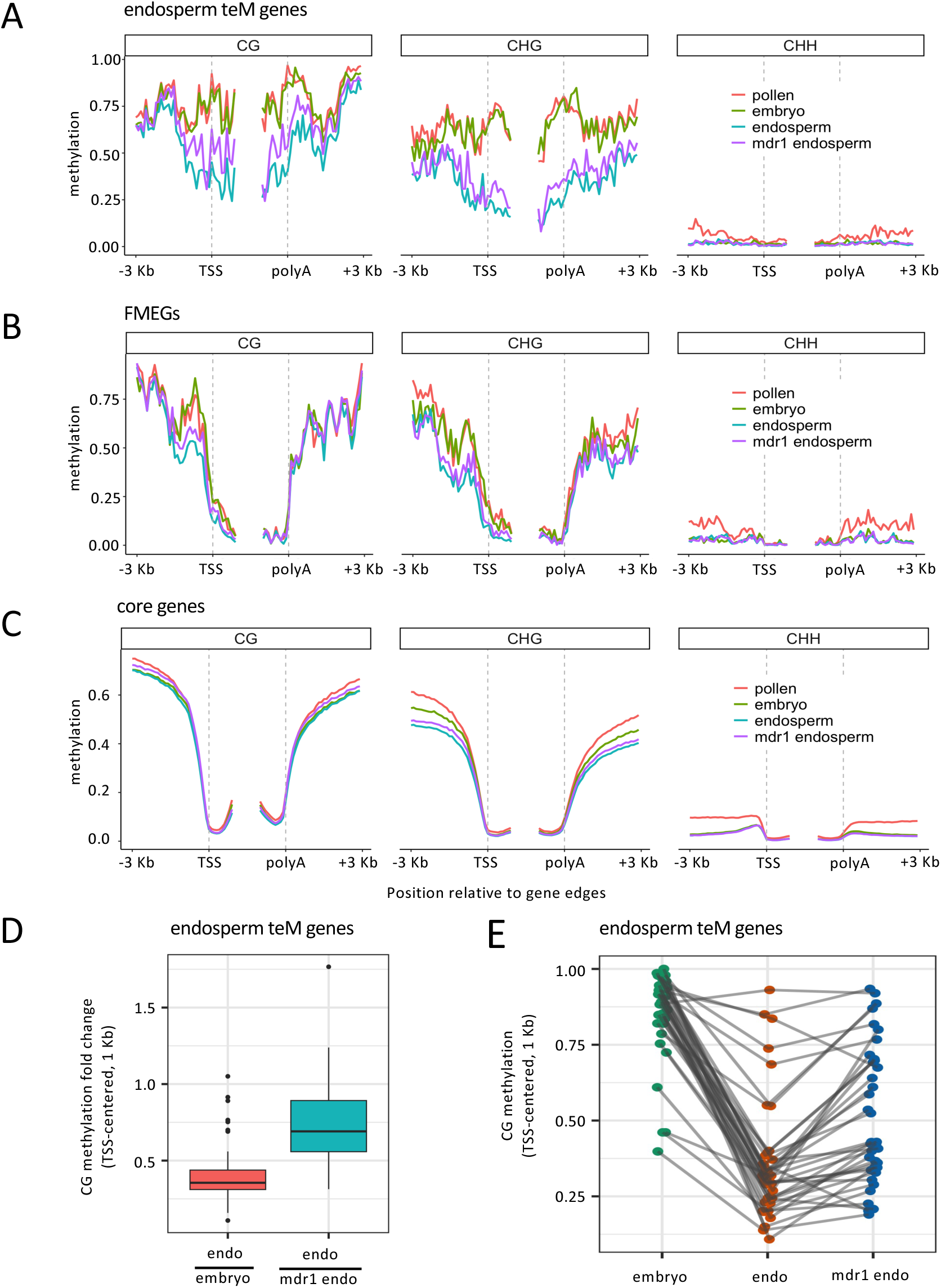
Endosperm teM genes have reduced CG and CHG methylation in endosperm. **a.** Metagene methylation profile for endosperm teM genes in the W22 genome. These are the 42 of 67 genes present in both the W22 and B73 genome annotations. Profiles are centered on transcription start sites (TSS) and polyadenylation sites (polyA). Methylation was measured on 100-bp intervals, from 3-Kb external to 1-Kb internal. **b.** Metagene methylation profile for the 54 FMEGs (flank-methylated endosperm genes) in the W22 genome. These represent classical imprinted genes that are maternally demethylated and expressed in endosperm. **c.** Metagene methylation profile for 28,291 core genes. Core genes are annotated at syntenic positions in W22, B73, and the 25 other NAM founder genomes. **d.** Methylation differences measured as fold changes. Boxplots show distributions of CG methylation for TSS-centered 1-Kb regions for each gene as fold changes between endosperm and embryo (both wildtype) or between wildtype endosperm and mdr1 mutant endosperm. **e.** CG methylation values underlying differences in (d).

Since many imprinted genes are regulated by DNGs, we asked how methylation patterns in endosperm teM genes compared with imprinted genes. Since imprinting can be mediated by polycomb repressive complex2 (PRC2) and histone methylation without TE-like DNA methylation (Batista and Köhler 2020), we created a list of high-confidence DNG-imprinted genes in maize by selecting genes with maternally preferred expression and which overlapped previously identified endosperm demethylated regions (DMRs) (Gent et al. 2022) in their gene bodies or in their 1-Kb flanking regions. We defined maternal preferred expression using published RNA-seq data from reciprocal W22 × B73 crosses (Higgins et al. 2025b). 61 core genes met these criteria. After removing seven which were also in our endosperm teM gene list, 54 were left. For this study, we refer to them as flank-methylated endosperm genes (FMEGs) because of their methylation and demethylation in flanking cis-regulatory elements regions, as is typical for imprinted genes. For clarity, in sporophyte, both FMEGs and endosperm teM genes can have methylation in their flanks (Fig. 2A-C). Both sets also have reduced methylation in their flanks in endosperm, especially in 5’ regions.

As an alternative means of quantifying methylation, we focused on the 1-Kb regions centered on transcription start sites and measured their average CG methylation for each gene. For this analysis, we included only genes which were methylated in embryo to allow for the possibility of detecting demethylation in endosperm. 37 of the 42 TSS-centered regions with CG methylation measurements had values above our cutoff value of 20% in embryo. Of these 37, 36 had decreased CG methylation in endosperm compared to embryo, with a median value in endosperm being 35.5% of the value in embryo (Fig. 2E,F). In theory, demethylation could be a passive outcome of dilution of methylated DNA with unmethylated DNA during replication rather than active demethylation by DNGs. The best test of this would be to compare wild-type with *mdr1 dng102* double mutant endosperm; however, double mutants fail early in development and are difficult to work with (Gent et al. 2022). Instead we used the *mdr1* single mutant which is fully viable but has a detectable increase in methylation at DNG target loci in endosperm. In spite of the noise introduced by random sampling of demethylated maternal reads and methylated paternal reads, 33 of 37 of the 1-Kb regions had decreased methylation in wild-type relative to *mdr1* mutant, with a median wild-type value being 69.1% of mutant endosperm (Fig. 2E F). Together these data reveal that like many imprinted genes, endosperm teM genes are demethylated by DNGs. Unlike most imprinted genes, however, they are demethylated not just in flanking cis-regulatory elements, but also throughout their coding DNA.

### High and endosperm specific expression of teM genes

By definition, endosperm teM genes had a minimum of fivefold more expression in endosperm than in the other nine tissues examined. However, the median expression difference was much higher, 258-fold, and the mean of 661-fold, indicating these genes were exceptionally endosperm specific (Fig. 3A,B). As a comparison we defined a set of endosperm genes using the same criteria as endosperm teM genes but without regard to methylation. Of these 1091 endosperm genes, which include the 67 teM genes, the median difference in expression was 12-fold, and the mean 132-fold. FMEGs had a slight preference for endosperm expression, median 1.1-fold, and a mean of 69-fold. Like their counterparts in pollen, the majority (53 of 67) of endosperm teM genes were encoded in two or fewer exons, 28 of these in a single exon (Fig. 3C). They also encoded short proteins with a median CDS length of 465 bp, compared to 1278 bp for FMEGs and 1056 bp for core genes (Fig. 3D). Endosperm teM genes were also among the highest expressing genes in endosperm. These 67 genes made up more than 40% of the transcriptome at 16 DAP (Fig. 3E). At every stage of development from 6-DAP to 34-DAP, at least 20 were in the top 1%, and at least 26 in the top 10% (Supplemental Fig. S1). At 8 DAP, 46 of 67 were in the top 10%.

**Fig. 3:**
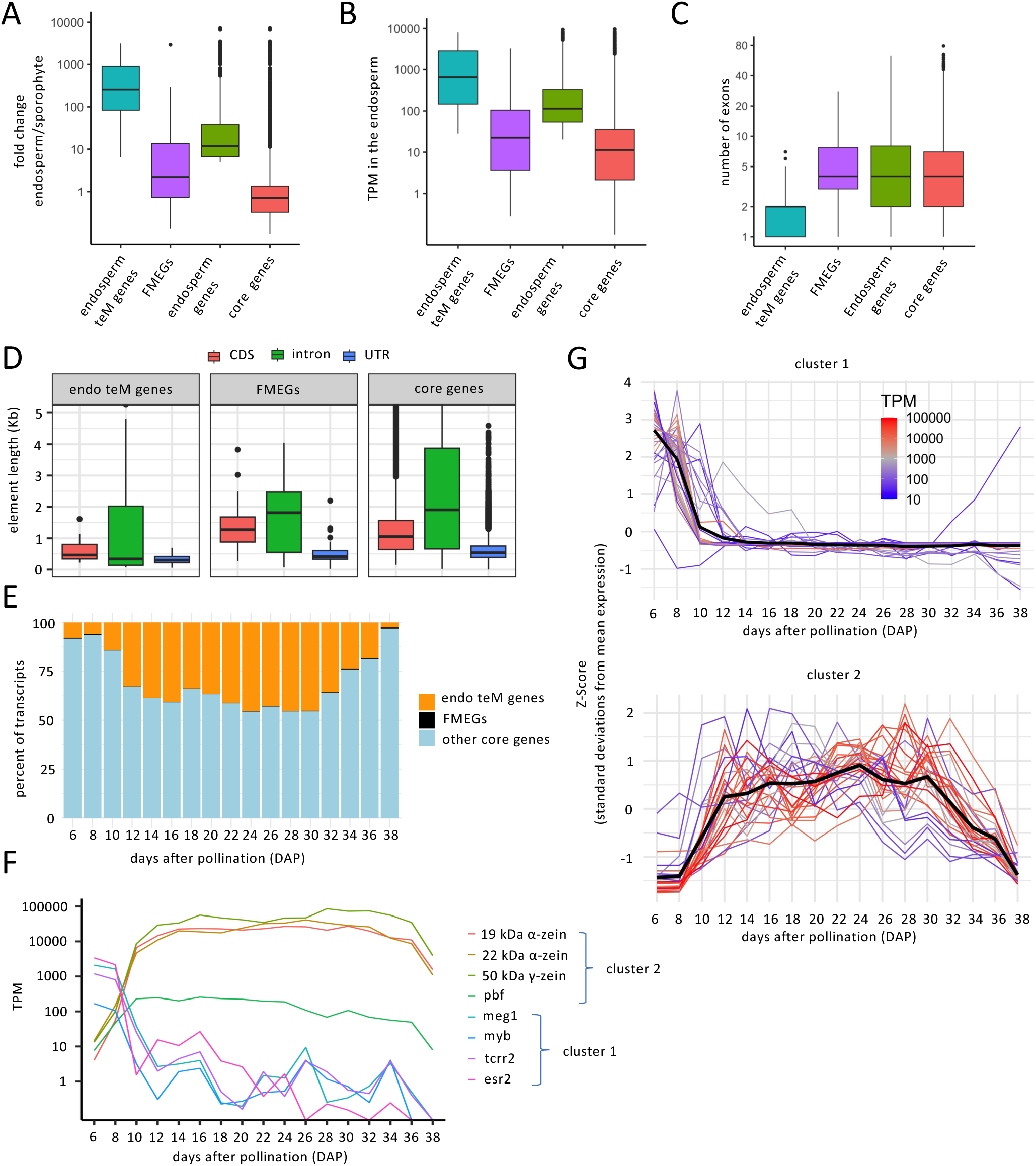
Expression patterns and structural features of endosperm teM genes. **a.** Endosperm teM genes are exceptionally endosperm specific. The Y-axis values indicate the ratio of the highest expression in endosperm development divided by the highest expression in any of the nine sporophyte transcriptomes from the NAM founder study. Box plots show the distribution for the 67 endosperm teM genes, the 54 FMEGs, and the 28,291 core genes (which include the other gene categories). Also included is a set of 1091 endosperm-specific genes that were defined with the same expression criteria as endosperm teM genes, but not constraints on methylation. These include the 67 teM genes. **b.** Endosperm teM genes are exceptionally highly expressed. Box plots show the distribution of maximum transcript per million (TPM) values for each gene over the endosperm development time course. **c.** Endosperm teM genes have few exons. **d.** Endosperm teM genes encode short proteins. **e.** Endosperm teM genes make up a large fraction of the endosperm transcriptome. **f.** Endosperm teM genes fall into two major clusters based on developmental expression profiles. The thick black lines indicate the average of all individual genes. **g.** Expression of example endosperm teM genes during development.

### TE-like methylation of secreted protein and seed storage protein genes

Endosperm consists of diverse cell types with different functions, and it undergoes major changes as it progresses from the central cell gamete, to the multinucleate coenocyte, to a predominantly starchy structure with few living cells (Wu et al. 2022). In rice and Arabiopsis, demethylation initiates in the central cell (Park et al. 2016). In maize, the fact that the *mdr1* mutant has an *r1* pigmentation phenotype in *mdr1* heterozygote endosperm when *mdr1* is maternally inherited but not paternally inherited also suggests a requirement in the central cell prior to inheritance of a functional paternal *mdr1* copy (Kermicle 1995, 1978). The fact that the *r1* pigmentation gene that is demethylated by DNGs is expressed in the outermost cell layer late in endosperm development suggests that the unmethylated state from the central cells can persist through endosperm development. To test whether teM genes show any characteristic expression patterns, we first clustered them using the same time course expression data that we used to identify them (Chen et al. 2014). This revealed two major clusters of genes, cluster 1 with 39 genes that peaked within the first eight days after pollination (DAP) and declined after, and cluster 2 with 28 genes that started low, increased rapidly at 8 DAP, and stayed high until 30 DAP (Fig. 3G,F and Supplemental Fig. S2). (Note that expression changes can reflect changes in the abundance of specific cell types over time, not just transcriptional regulation within cells). FMEGs produced a more variable set of expression patterns than teM genes and did not produce clear clusters (Supplemental Fig. S2).

Cluster 1 of the endosperm teM genes had wide-ranging spatial expression patterns and gene functions. Some genes, such as the carboxypepsidase Zm00001eb240770 or the ubiquitin paralogs Zm00001eb240770 and Zm00001eb275020 are expressed across endosperm compartments (Zhan et al. 2015). There was a trend among Cluster 1 genes, however, for expression in the peripheral cell layers of the endosperm and secreted proteins. These layers, called the embryo surrounding region (ESR), basal endosperm transfer layer (BETL), and aleurone, are important for signaling between endosperm and embryo or between endosperm and the maternal sporophyte (Wu et al. 2022). Sixteen of the 39 cluster 1 genes were clearly in this group: the defensin-like BETL1/BET1 (Gómez et al. 2009), two tandem duplicates of the antifungal protein BETL2/BAP1 (Serna et al. 2001); three lipid transfer proteins, BETL9/NPTD6, AL9, and PLT32/NPTD16 (Royo et al. 2014; Fang et al. 2023; Zhan et al. 2015); five tandem duplicates of the BETL developmental regulator MEG1 (Costa et al. 2012); the protein of unknown function AE1/ANE1 (Magnard et al. 2000); the endopeptidase EBE2 (Magnard et al. 2003); the CLAVATA3/ESR homolog ESR2 (Bonello et al. 2002), and two two-component response regulators, TCRR1 and TCRR2 (Muñiz et al. 2010). The transcription factor MYBR19 that activates BETL genes was also in cluster 1 (Yuan et al. 2024).

Zeins and closely related prolamins encode signal peptides and behave like secreted proteins in that they have signal peptides and localize to the endoplasmic reticulum (ER). But rather than being secreted, they can polymerize in the ER to form protein storage bodies that make up as much as 70% of the total protein in endosperm, the majority of which is α-zein (Holding 2014). The less abundant β- and γ-zein proteins are broadly conserved in grasses, in the same group as glutelins in wheat (where they are a component of gluten). The other type of zein proteins, δ-zeins, are more similar to α-zeins and together make a group that is common to the C4 panicoid grasses (e.g., maize, sorghum, sugarcane) (Wu and Messing 2012; Sabelli and Larkins 2009; Holding 2014). 22 of 28 cluster 2 genes are α-zeins (fourteen 19-kDA α-zein, and eight 22-kDA α-zeins), and one is the single maize 50-kDa γ-zein. Cluster 2 also includes *pbf1*, a transcription factor that promotes zein expression (Zhang et al. 2016). Unlike cluster 1 genes with predominant expression in peripheral cell layers, zeins express in the starchy endosperm. In midstage endosperm development, α-zeins make up about 50% of the total endosperm transcripts (Chen et al. 2014). Maternal demethylation of α-zein genes has been detected previously, but causal relationships between methylation, demethylation, and transcription have been difficult to define (Lund et al. 1995; Wu and Messing 2012). α-zein genes are short and encoded in multiple copies which can complicate mapping of RNA-seq reads to any single gene. Thus, we counted how many annotated zein genes were categorized as teM genes, regardless of expression measurements (Zeng et al. 2023). All eighteen of the 19-kDa α-zeins, eleven of the 22-kDa α-zein genes, and the single 50-kDa γ-zein genes were teM genes (Supplemental Dataset 2). The single 16-kDa and 27-kDa γ-zeins, the single 10-kDa δ-zein, and the single 15-kDa β-zein genes were unmethylated; and the single 18-kDa δ-zein had intermediate methylation. It is challenging to determine whether the demethylation of prolamins is a conserved process because they have high copy number variation, high substitution rates, and lack synteny between species (Xu et al. 2011). However, in Sorghum, methylation-sensitive restriction digests suggest that α-kafirin genes–homologous to α-zeins–are methylated in leaf and demethylated in endosperm (Bedell et al. 2005; Zhang et al. 2007). More distant conservation of this phenomenon is supported by the identification of several glutelin genes among demethylated regions in rice endosperm and downregulation of glutelins in a rice DNG mutant (Irshad et al. 2022; Zemach et al. 2010).

### Acquisition of both active and repressive chromatin modifications with demethylation

DNA methylation is part of multiple layers of chromatin-based constitutive silencing of TEs mediated by proteins with methyl binding domains (Boone et al. 2023). The dynamic repression of genes in response to developmental or environmental cues in plants typically relies on histone modifications, histone variants, and other chromatin-bound factors that do not include methyl binding domains. A prediction of endosperm teM genes is that their TE-like DNA methylation in sporophyte would correspond to TE-like histone modifications and exclusion of ones associated with dynamic gene regulation. Consistent with this, analysis of both our own histone modification data and published data revealed a striking absence of genic histone modifications (Fig. 4 and Supplemental Fig. 3) (Ricci et al. 2019; Cahn et al. 2024). Also absent from teM genes were accessible chromatin regions and H2A.Z. In contrast, they had heterochromatic modifications H3K27me2 and H3K9me2 profiles similar to TEs and unlike genes (Gent et al. 2014). Chromatin of pollen teM genes was similar to endosperm teM genes.

**Fig. 4:**
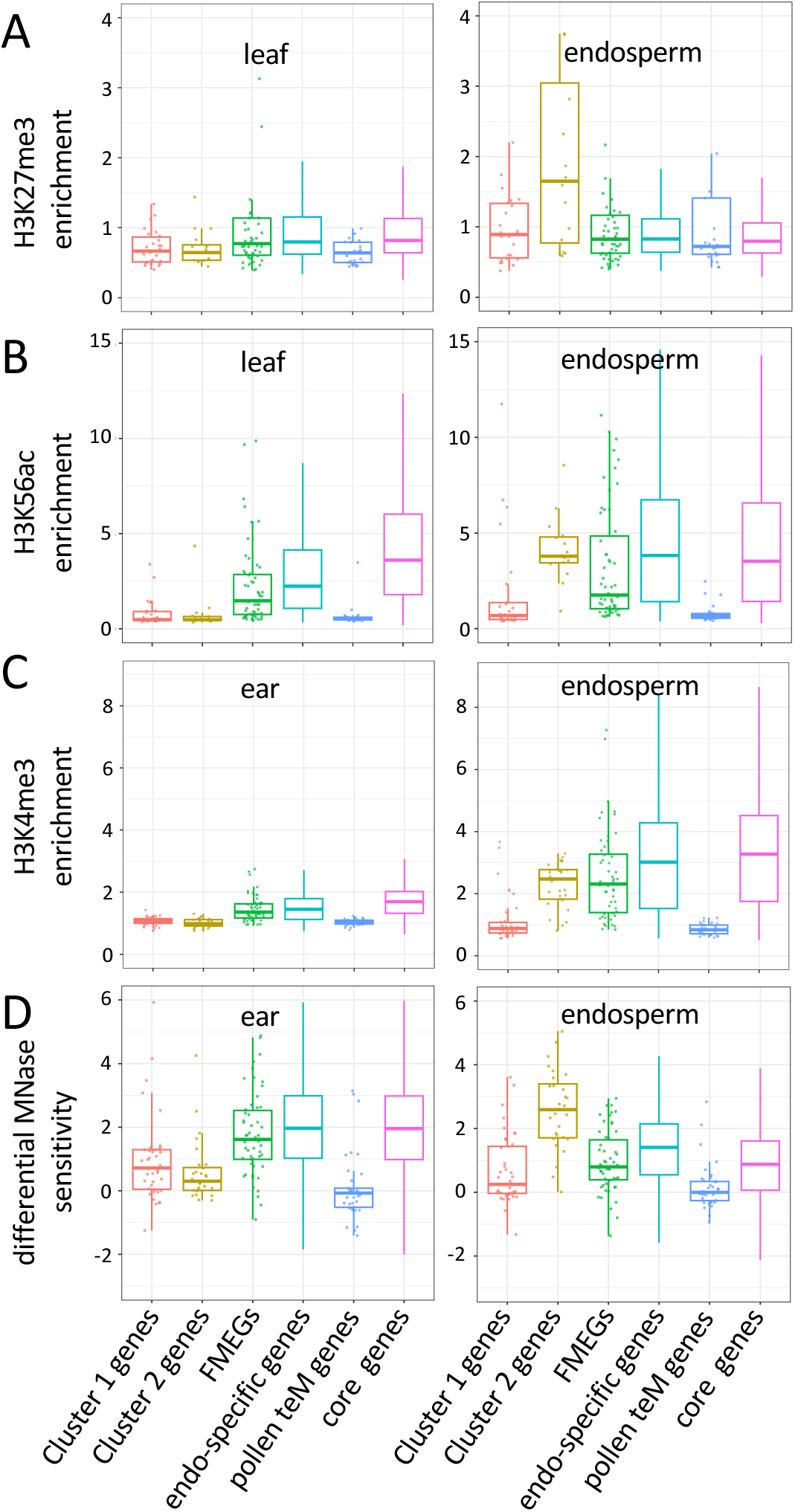
TE-like chromatin modifications of teM genes in sporophyte, gene-like in endosperm. **a-b.** CUT&Tag with antibodies against H3K27me3 or H3K56ac show enrichment in endosperm in 2-KB regions centered on transcription start sites (TSSs) of cluster 2 genes. Enrichment is normalized read count from CUT&Tag with indicated antibodies normalized by CUT&Tag with IgG. Endosperm is from 15 days after pollination. Data is from W22. **c** ChIP with antibodies against H3K4me3 show enrichment in endosperm in 2-KB regions centered on TSSs of cluster 2 genes. Enrichment is normalized read count from ChIP with H3K4me3 antibodies normalized by reads from sequenced input. Data is from B73, from Cahn et al (2024). **d** Differential micrococcal nuclease (MNase) sensitivity assays show accessibility in endosperm in 2-KB regions centered on TSSs of cluster 2 genes. Positive values indicate MNase hypersensitivity, whereas negative values indicate MNase-resistant chromatin. Data is from B73, from Cahn et al (2024).

Given that endosperm is a complex tissue with cell-type-specific gene expression patterns, it seemed likely that some of the demethylated genes would have histone modifications that repress gene expression. To test this we generated histone modification profiles for H3K27me3, a product of PRC2, and for one representative active chromatin modification in endosperm, H3K56 acetylation (H3K56ac). At 15 DAP, cluster 2 endosperm teM genes had both modifications, as well as H3K4me3 and accessible chromatin (Fig. 4). We expect the presence of both active and repressive chromatin signatures is a result of different activity states in different endosperm cell types. Consistent with their H3K27me3 in endosperm, PRC2 mutants in maize have elevated zein expression (Wang et al. 2025a). These results of TE-like chromatin in sporophyte but gene-like in endosperm are consistent with constitutive, robust silencing throughout the plant life cycle from generation to generation but dynamic gene regulation in endosperm.

### Methylation in promoters, not gene bodies linked to imprinting of endosperm teM genes

Specific alleles of α-zein genes have maternal preferred expression (Lund et al. 1995; Higgins et al. 2025b), as does a *meg1* gene (Gutiérrez-Marcos et al. 2004). Quantifying imprinting relies on use of hybrid endosperm with genetic variants to distinguish between parental alleles. This becomes problematic for short genes, especially ones with multiple copies, where RNA-seq reads do not clearly distinguish between gene copies. Thus absence of evidence for imprinting from existing studies may not indicate absence of imprinting. To more thoroughly investigate the extent of imprinting among the endosperm teM genes, we examined the source data from a recent study that included eight different genetic backgrounds and multiple timepoints (Higgins et al. 2025b). Because of the variability of imprinting between stages of endosperm development and between alleles of the same gene, we only considered genes with consistent maternal expression ratios across multiple genotypes. 13 of 67 endosperm teM genes passed our quality control standards for consistency (Supplemental Dataset 1). Without imprinting, the expected maternal ratio of expression is ⅔ becuase endosperm carries two maternal genomes for each paternal genome. With threefold maternal preference, the maternal ratio would be 6/7 (∼0.86). Six of the 13 exceeded that threshold (Fig. 5A). In comparison, 51 of 10,700 qualifying core genes did. This amounts to a 97-fold enrichment for maternal preference among endosperm teM genes.

**Fig. 5:**
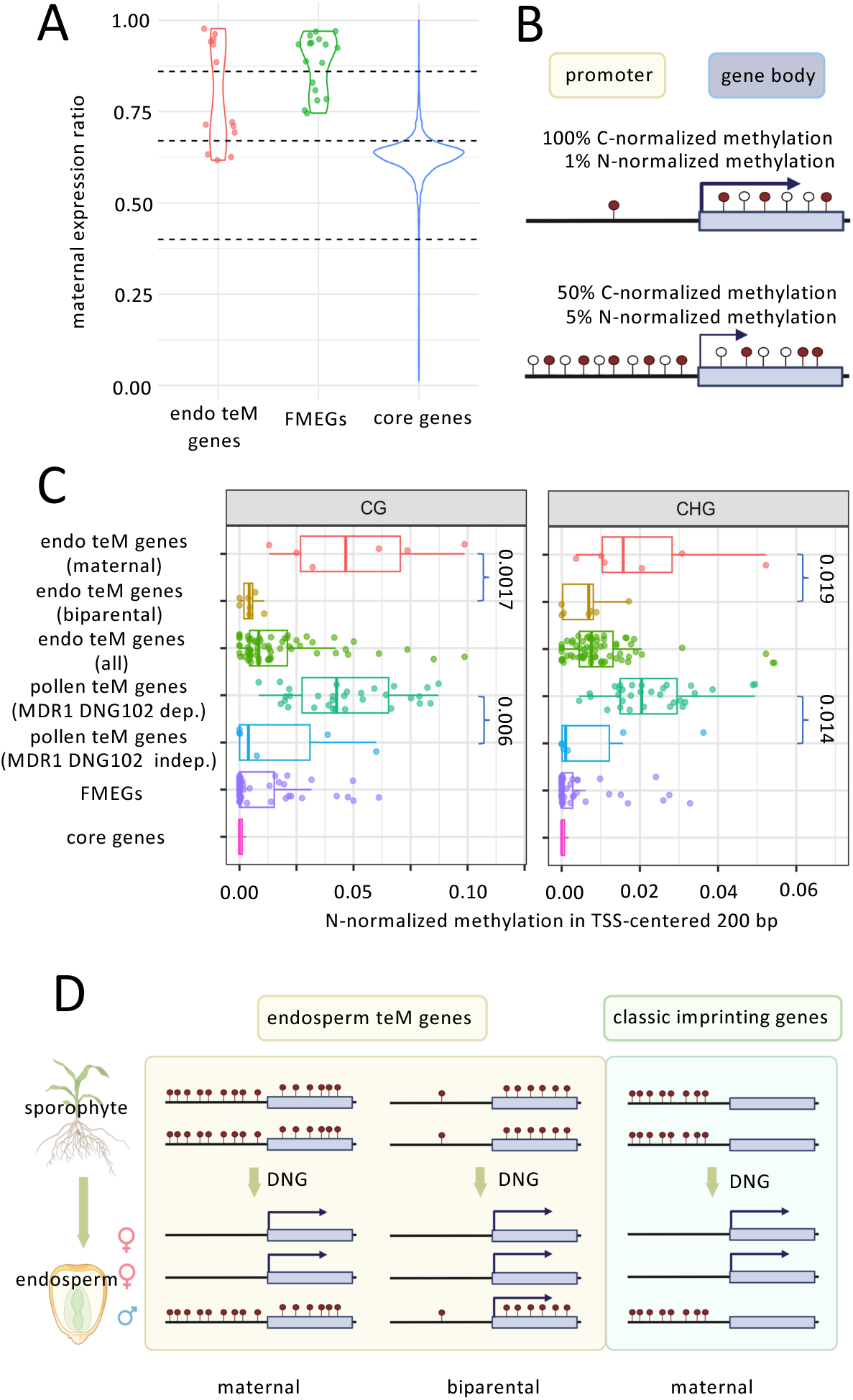
Imprinting of endosperm teM genes correlates with promoter methylation. **a.** Some endosperm teM genes have imprinted expression, some do not. Maternal expression ratio is the proportion of maternally-derived transcripts as measured by parent-specific sequence variants in RNA-seq data from hybrid endosperm. The upper dashed line indicates a maternal expression ratio of 6/7 (threefold more than expected), the middle line indicates a ratio of 2/3 (expected), and the lower line indicates a value of 2/5 (threefold less than expected). **b.** A schematic of nucleotide-normalized (N-normalized) methylation vs cytosine-normalized methylation (C-normalized). Open lollipops represent unmethylated cytosines, solid lollipops represent methylated cytosines. **c.** N-normalized methylation in the 200-bp region centered on transcription start sites (TSSs) of endosperm teM genes correlates with dependence on demethylation for expression. For endosperm teM genes, “maternal” are the six genes in (a) with maternal expression more than threefold greater than expected, “biparental” are the seven with expected maternal expression, and “all” is the complete set of 67. For pollen teM genes, “MDR1 DNG102 dep” are the set of 31 whose pollen expression depends on these two DNGs, and “MDR1 DNG102 indep” are the set of six whose pollen expression is independent of them. **d.** A model for imprinting of endosperm teM genes based on methylation in promoters.

The lack of maternal preference for seven of the measured genes indicates either little or no impact of paternal DNA methylation on their expression in endosperm, or that paternal copies are also demethylated. To test this, we examined parent-specific methylation data from B73/W22 hybrid endosperm (Higgins et al. 2025a). While there is a global and biparental reduction in CHG methylation in endosperm (Fig. 2B), the change in 5’ regions of endosperm teM genes was relatively minor: paternal CHG methylation of 51% in mature endosperm compared to 74% in leaf (Supplemental Fig. S5). A plausible explanation for paternal expression of endosperm teM genes in endosperm but not in sporophyte is that a partial loss of methylation combined with the presence of endosperm specific TFs such as O2 and PBF1 (Wu et al. 2022) allows for remodeling of heterochromatin into euchromatin and associated high expression for some but not others.

The others, with maternal preferred expression–e.g., *meg1* genes, some zeins, and the six of 13 identified here–present an interesting problem. What distinguishes them from the biparentally expressed ones? Given the stronger connections between methylation in cis regulatory elements and gene silencing than between between methylation in gene bodies and gene silencing (Zeng et al. 2023; Hufford et al. 2021), we hypothesized that the amount of methylation in promoters in endosperm teM genes would inversely correlate with paternal expression. CG methylation in particular should have a strong effect since methylation in this context often reaches 100% (Supplementary Fig. S5). To test this, we took into account not only methylation in the conventional way, as the proportion of methylated cytosines to total cytosines at each position, but also considered the number of cytosines in each promoter. The rationale behind this is five methylated cytosines in a 100 bp region with ten total cytosines would normally have a greater effect on chromatin structure than one methylated cytosine would, even if it were the only cytosine in the region (Fig. 5B). Normalizing by the total number of nucleotides (N) rather than just the total number of cytosines would give values of 5% for the first case but 1% for the second. Consistent with our hypothesis, the six genes with strong maternal preference had a mean N-normalized CG methylation value of 5.1%, while the seven without maternal preference had a mean N-normalized CG methylation value of 0.44% (Fig. 4C and Supplemental Fig. S6). This difference was statistically significant (p-value = 0.0017, one-tailed Wilcoxon rank-sum test). There was also a significant difference in promoter CHG methylation even though CHG methylation is globally reduced in endosperm (2.1% vs 0.58%, p-value 0.019). Methylation levels in gene bodies did not correlate with imprinting (Supplemental Fig. S6). Extrapolating from the 13 tested endosperm genes, we expect based on their promoter methylation levels in B73 leaf that about 30 of 67 would also be maternally preferred (Fig 5C). Eight of these were zeins (Supplemental Dataset 1).

### Dependence of pollen teM genes on MDR1 and DNG102 linked to methylation in promoters

While the majority of pollen teM genes (called methylated pollen genes in the prior study) are strongly downregulated in *mdr1 dng102* double mutant pollen, some retain high expression (Zeng et al. 2024). Of 56 pollen teM genes identified, 37 had high confidence expression measurements in the double mutants, with 31 being called as differentially expressed and six retaining high expression. We had hypothesized that one or both of the other DNGs encoded in maize (DNG103 and DNG105) might demethylate these six genes in pollen. However, our finding that methylation in promoters of endosperm teM genes associated with paternal silencing led us to ask whether the six pollen genes that appeared to be unaffected in *mdr1 dng102* double mutants might simply have fewer cytosines in their promoters. Supporting this hypothesis, the differentially expressed pollen teM genes had a mean N-normalized CG methylation value of 4.7% in promoters while the unchanged genes had a value of 1.8% (with p-value 0.006) (Fig. 5C). CHG methylation in their promoters also strongly correlated with differential expression, with mean N-normalized CHG methylation value of 2.3% for differential expressed genes and 0.9% for unchanged genes (with p-value 0.014). Together with the results from the 13 endosperm teM genes that have high-confidence measures of imprinted expression in endosperm, these results suggest that methylation near transcription start sites rather than in the gene body is key to silencing teM genes, at least in endosperm and pollen cells which are primed to express these genes.

## DISCUSSION

While endosperm teM genes and classical imprinted genes share a mode of gene regulation–demethylation of upstream elements by DNGs–teM genes have several major distinctions. First, teM genes have few introns and encode short proteins. Second, they are exceptionally tissue specific and highly expressed. Third, their chromatin profiles in the sporophyte look like TEs. Lastly, their methylation is throughout their gene bodies. Unlike CG gene body methylation, which is associated with constitutive gene expression in sporophyte and endosperm, TE-like methylation in gene bodies is clearly associated with repression (Zeng et al. 2023) (Fig. 6). How TE-like DNA methylation functions in gene regulation has been interpreted largely through the lens of genes like FWA in Arabidopsis, where methylation of TE fragments in its promoter prevents ectopic expression outside of endosperm (Kinoshita et al. 2007). Many other related examples have also been reported where methylation of TEs and other repetitive DNA in cis regulatory elements makes gene expression responsive to DNA demethylation, e.g, *ROS1* and *SDC* in Arabidopsis; and *b1*, *r1*, *pl1*, and *ufo1* in maize (Sidorenko et al. 2024; Williams et al. 2015; Henderson and Jacobsen 2008; Belele et al. 2013; Walker 1998; Deans et al. 2024; Wittmeyer et al. 2018). In contrast to these unusual situations, gene regulation in plants does not normally involve DNA demethylation because cis-regulatory elements are constitutively unmethylated (Crisp et al. 2020). For some cis-regulatory elements, that may require surveillance by DNGs to actively remove methylation (Wang et al. 2025b; Halter et al. 2021). In contrast, teM genes that are highly expressed in pollen and endosperm are special cases where DNA methylation and demethylation directly regulate gene expression.

**Fig. 6:**
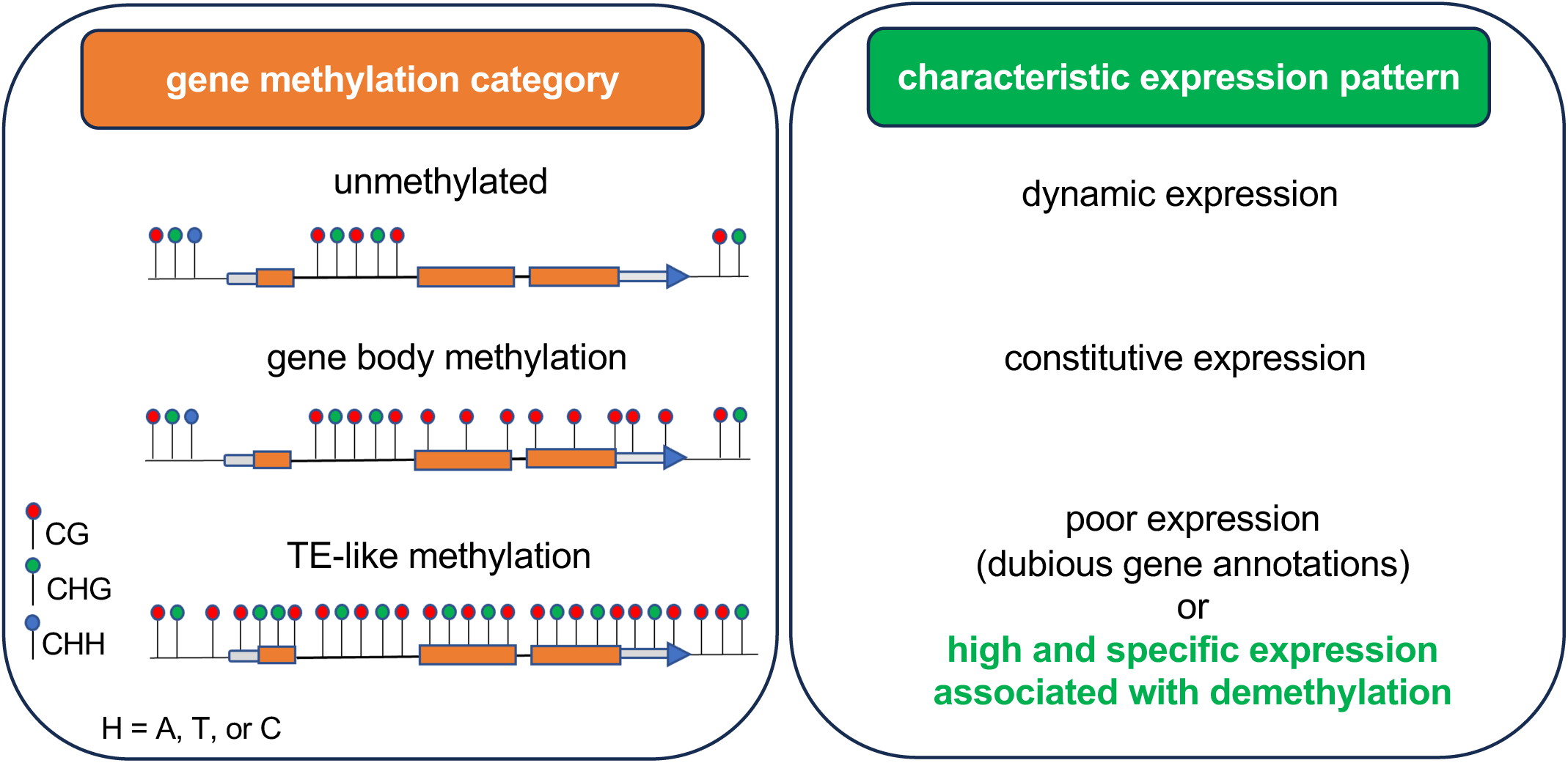
Schematic of gene methylation categories. While the vast majority of genes with TE-like methylation may reflect misannotated TEs or pseudogenes, a few are highly expressed and highly specific and function in pollen and endosperm. Figure adapted from Zeng et al. 2023 <u>^41^</u>.

A shared pattern in pollen and endosperm teM genes is their encoding proteins enter the secretory pathway. In pollen, their destination is the apoplast, where they can modify cell walls (Zeng et al. 2024). In endosperm, many are transferred between cells. Most notable, however, are the zeins, which enter the secretory pathway but stay in the ER to form protein bodies that are characteristic of endosperm of maize and related grasses (Holding 2014; Wu and Messing 2012). The diversity of endosperm teM genes, including ones with likely cytoplasmic or nuclear functions such as transcription factors and ubiquitin components suggests roles in a broad range of processes. Their being broadly demethylated both spatially and temporally in endosperm raises the question of how they have specific expression patterns within endosperm (Fig. 3). Part of the answer is that lack of methylation is not sufficient for transcriptional activation. As in sporophyte, the majority of genes are unmethylated in endosperm regardless of expression (Fig. 2). Endosperm teM genes still require transcription factors even in their demethylated state (Zhang et al. 2016). The dynamic levels of H3K27me3 of endosperm teM genes in endosperm specifically suggest that their expression patterns are also regulated by PRC2 when they are demethylated (Fig. 4).

In pollen, highly cell-type-specific and strong expression of most but not all teM genes is directly dependent on demethylation by MDR1 or DNG102 (Zeng et al. 2024). Some endosperm teM genes behave similarly, with high expression and specificity limited to the demethylated maternal genome alleles. Some, however, have high specificity and expression from both parental genomes in spite of the paternal alleles retaining CG methylation at sporophyte levels. We found that the difference between the maternal preferred teM genes and biparental ones could be explained by methylation in promoters rather than in gene bodies. Supporting this, we found that dependence of pollen teM genes on MDR1 and DNG102 also correlates with methylation in promoters (Fig. 5). This raises the question of why some genes would have TE-like methylation in just their gene bodies and not in promoters. One possibility is that methylation in gene bodies has a repressive function in the sporophyte. The other is that the methylation in gene bodies is not functional and arises as a consequence of the genes’ never being expressed in the sporophyte lineage. In that case, TE-like methylation in gene bodies would be a signature of repression in sporophyte rather than a cause. What is clear is that removal of TE-like methylation, most crucially near transcription start sites, is essential for expression of some of these genes, as evidenced both by strong maternal expression preference in endosperm and by the loss of expression in DNG mutant pollen (Zeng et al. 2023). It is also clear that the genes regulated in this way are exceptionally highly expressed and encode functional and conserved proteins, including many secreted proteins and seed storage proteins.

## METHODS

### Identification of endosperm teM genes and other gene sets from expression and methylation data

Using the Zm-B73-REFERENCE-NAM-5.0 annotation, we defined core genes present in all 25 NAM genome lines and B73 as previously described (Hufford et al. 2021) based on the pan_gene_matrix_v3_cyverse.csv downloaded from https://de.cyverse.org/anon-files//iplant/home/shared/NAM/NAM_genome_and_annotation_Jan2021_release/SUPPLEMENTAL_DATA/pangene-files/pan_gene_matrix_v3_cyverse.csv (this data has since then been removed from Cyverse but is now available at https://github.com/dawelab/endo-teM-genes/blob/main/identify_imprinting/pan_gene_matrix_v3_cyverse.csv.zip). This produced a total of 28,291 core genes. To identify endosperm teM genes, we collected the B73 RNA-seq data from a previous study (Chen et al. 2014) spanning 6 to 38 days after pollination (DAP). All 21 samples were downloaded directly from Sequence Read Archive (SRA) under accession number SRP037559 using SRA-Toolkit v3.0.3 and parallel-fastq-dump v0.6.7. The raw sequences were trimmed with Trim Galore v0.6.7 with the parameter --paired (https://github.com/FelixKrueger/TrimGalore). Then the trimmed reads were aligned to the B73 v5 genome using STAR v2.7.10b (Dobin et al. 2013). Prior to alignment, a STAR index was generated with the parameter --sjdbGTFtagExonParentTranscript Parent to format the GFF3 annotation file. STAR mapping was performed with default parameters. Subread v2.0.6 (Liao et al. 2013) was used to count the paired-end reads of each transcript within the exons using -p --countReadPairs -t exon -O -g Parent. The -O -g Parent options ensured that the reads mapped to multiple transcript isoforms were properly assigned. Maize reference genome sequence and annotation data were downloaded from https://download.maizegdb.org/Zm-B73-REFERENCE-NAM-5.0/. The resulting raw read counts were filtered to retain only canonical transcripts, then expression values averaged across biological replicates. Gene expression was normalized using the TPM (Transcripts Per Million) method to account for sequencing depth and transcript lengths. Methylation data from developing second leaves and TPM values from nine sporophyte tissue samples were obtained from prior work (Zeng et al. 2023) which was derived from the NAM founder genome assembly project(Hufford et al. 2021) (https://raw.githubusercontent.com/dawelab/Natural-methylation-epialleles-correlate-with-gene-expression-in-maize/refs/heads/main/Data/B73.all.csv). The nine tissue samples were embryo, tassel, anther, shoot, ear, leaf tip, leaf middle, leaf base, and root.

Genes were classified as teM genes if they had at least ten informative CHGs (ones that were spanned by EM-seq reads) and methylation values of at least 40% for CG and for CHG in leaf. Here and elsewhere, methylation values were defined using the conventional C-normalized method unless stated otherwise. To select for teM genes that are enriched for endosperm expression, we required a ≥ 5-fold increase in TPM in at least one stage of endosperm development compared to all other nine sporophyte samples. In addition, a minimum TPM of 20 in at least one endosperm stage was required to filter out low-expressed genes with unreliable TPM fold change measurements. This produced a set of 119 genes. After removing ones that were not in the core gene set, this analysis identified 67 endosperm teM genes. To define a control set of genes, all the same criteria were used except without regard to DNA methylation. This produced a set of 1,091 genes that we refer to as endosperm-specific genes.

To generate a representative set of DNG-dependent classical imprinted genes, we used allele-specific expression data from a recent study where RNA-seq from reciprocally crossed W22 and B73 endosperm at 11, 14, 17, and 21 DAP was used to calculate the maternal allele expression ratio based on sequence variants associated with each genome (Higgins et al. 2025b). We filtered out the genes with RPM < 1 at all four time points to avoid misleading maternal preference caused by extremely low expression. Genes with maternal ratios of ≥ 0.9 were classified as MEGs; genes with maternal ratios of ≤ 0.3 were PEGs. Genes that met criteria for both MEG and PEG at different time points were excluded. Gene ID conversion between B73 and W22 annotations were conducted based on MaizeGDB_maize_pangene_2020_08.tsv downloaded from https://download.maizegdb.org/Pan-genes/archive/pan-zea-2020/. This identified 295 core genes which were MEGs. To identify the subset of imprinted genes regulated by demethylation in endosperm, we selected all that overlapped with endosperm demethylated methylated regions (DMRs) anywhere in their gene bodies or 1-Kb flanking regions. These DMRs were identified previously through a comparison between endosperm and embryo (Gent et al. 2022). After removing seven genes from this list that were in our endosperm teM genes and Z that were PEGs, this produced a list of 54 maternally expressed imprinted genes that we call flank-methylated endosperm genes (FMEGs).

### K-means clustering of expression patterns

To characterize expression patterns and cluster genes accordingly, we calculated Z-scores for the TPM values across 6 to 38 DAP using the formula z = (x - μ) / σ, where μ and σ are the mean and standard deviation of TPM for each gene. The factoextra 1.0.7 R package (https://github.com/kassambara/factoextra) was used to determine the optimal number of clusters using both the Elbow Method and Silhouette Method, both of which indicated k=2 is optimal (Supplemental Fig. S2). Then the kmeans R function was used to cluster the 67 endosperm teM genes into 2 groups. Gene names, protein functions, and families were manually curated using data from MaizeGDB (Portwood et al. 2019) and by blasting protein sequences to identify conserved domains. Figures were generated by ggplot2 v3.5.1 (Wickham 2016).

### Imprinted expression analysis

Since the teM endosperm genes tend to have fewer exons and be encoded in multicopy gene families, making them more likely to have multimapping issues, we applied a stringent criterion to assess their imprinting status. We obtained a table of expression values, “mat_pref_NAM.txt” from a study in which B73 was reciprocally crossed with 8 inbred lines: W22, B97, Oh43, Ky21, NC358, CML333, M162W, and Ki11 (Higgins et al. 2025b). Gene IDs were converted to B73 annotation using MaizeGDB_maize_pangene_2020_08.tsv downloaded from MaizeGDB at https://download.maizegdb.org/Pan-genes/archive/pan-zea-2020/. Genes for which at least one allele had ≥ 5 RPM in at least one cross and with maternal preference variance ≤ 0.02 across all crosses that met the ≥ 5 RPM threshold were selected for further analysis. This stringent method produced 11,806 genes with high-confidence imprinting data. The mean maternal preference values for the eight crosses were used to represent imprinting status.

### Identification of zein genes

We identified zeins genes in the maize genome by searching for specific keywords in MaizeGDB (Portwood et al. 2019) : Z1 for α zeins, bz15 for beta zeins, gz for gamma zeins, and dzs for delta zeins. from B73v5 with pan-genome IDs, a reference table linking gene IDs to pan IDs was retrieved from (https://raw.githubusercontent.com/dawelab/Natural-methylation-epialleles-correlate-with-gene-expression-in-maize/refs/heads/main/Data/B73.all.csv). Genes sharing the same pan ID as known zeins were incorporated into the analysis. Methylation information for these genes was obtained from the same data.

### CUT&Tag

We harvested endosperm samples from a W22 stock 15 days after pollination (DAP) when embryos were approximately 2 mm wide by 3.5 mm long (BioSample SAMN26561113). Endosperm was separated from pericarp, nucellus, and embryo by hand dissection with forceps. Leaf samples were harvested from a W22 stock at the stage prior to unfurling of 1st leaf (but excluding the first leaf itself), similar to the leaf samples used in the published ChIP experiments (BioSample SAMN26561117) (Ricci et al. 2019). After freezing in liquid nitrogen, nuclei were extracted and used for CUT&Tag using a CUT&Tag-It kit (Active Motif 53160) as described previously (Dawe et al. 2023). Three biological replicates with antibodies against H3K27me3 (Active Motif 39157) and H3K56ac (Active Motif 39282) were used as well as an IgG control (EpiCypher 13-0042). Nuclei containing 500-800 ng of DNA were used as input for each CUT&Tag reaction, and libraries were amplified using 13 or 14 cycles of PCR.

### CUT&Tag, ChIP-seq, and ATAC-seq and MNase-seq analyses

CUT&Tag libraries were Illumina sequenced paired-end with 150 nt read lengths. Reads were trimmed of adapters using Cutadapt (v2.8) (Martin 2011) with parameters -q 20 -a CTGTCTCTTATACACATCT -A CTGTCTCTTATACACATCT -O 1 -m 50 and mapped to either W22 (Springer et al. 2018) or B73 (Hufford et al. 2021) reference genomes using Bowtie2 (v2.5) (Langmead and Salzberg 2012). Alignments were filtered to retain uniquely mapped reads with MAPQ ≥ 20. The read coverages were generated with deepTools (v3.5.5) bamCoverage in 10bp bins and normalized to RPGC using --normalizeUsing RPGC –effectiveGenomeSize 2300000000 (Ramírez et al. 2014). Enrichment values were computed with deepTools bigwigCompare using CUT&Tag with IgG as the control. The metagene profiles centered on TSSs (transcription start sites) and polyadenylation sites were generated using deepTools computeMatrix using 10-bp bins.

For ChIP-seq, ATAC-seq (assay for transposase-accessible chromatin) and MNase-seq (micrococcal nuclease), reads were processed, mapped to the B73 reference genomes, normalized read coverages calculated, and metagene profiles produced using the same workflow as CUT&Tag except that adapter trimming and quality filtering were performed using Cutadapt (v4.9) (Martin 2011) with default settings, and PCR duplicates were removed using Picard MarkDuplicates (v3.3.0) ([CSL STYLE ERROR: reference with no printed form.]). Enrichment values were computed with deepTools bigwigCompare using either randomly fragmented DNA (for H3K9me2 and H3K27me2 ChIP-seq) or input control reads (all other ChIP-seq and for ATAC-seq) as in the original studies (Ricci et al. 2019; Cahn et al. 2024; Gent et al. 2014). For MNase-seq, the heavy-digestion normalized coverage was subtracted from the light-digestion coverage instead of computing enrichment relative to a control (Turpin et al. 2018). The H2A.Z, H3K27ac, H3K27me3, H3K36me3, H3K4me1, H3K4me3, and H3K56ac, H3K9ac CHIP-seq, and ATAC-seq were from developing second leaves (Ricci et al. 2019). H3K9me2 and H3K27me2 ChIP-seq was from multiple V4 stage developing leaves (Gent et al. 2014). H3K4me3 ChIP-seq was from both 15-DAP endosperm and immature ears (Cahn et al. 2024). MNase-seq was also from 15-DAP endosperm and immature ears (Turpin et al. 2018).

### DNA methylation analyses

Whole genome methylation CGmaps produced by previous studies were used as the data sources for measuring methylation values: developing 2nd leaf from B73 (Hufford et al. 2021), mature pollen from W22 (Zeng et al. 2024), 15-DAP endosperm (wild-type and *mdr1* mutant) and embryo from W22 (Gent et al. 2022), and mature endosperm and developing 2nd leaf from B73 X W22 hybrids (Higgins et al. 2025a). The CGmapTools (Guo et al. 2018) v1.3 MTR tool was used to measure methylation over individual loci using the “by site” method. Metagene methylation measurements were obtained using the CGmapTools MFG tool on 100 bp intervals. For N-normalized methylation values, the mC/total C methylation values for each region was multiplied by the number of cytosines (context specific) and divided by the total number of nucleotides. To quantify cytosines in specific sequence contexts, we implemented a custom function named count_cytosines in R. For each gene sequence, both the forward strand and reverse complement were considered. Counts were obtained by sliding a window of 2 or 3 nucleotides across the sequence to identify CG, CHG, and CHH sites. Cytosine contexts were counted if the cytosine itself resided within the analyzed region. Parent-specific methylation values were calculated using the CGmapTools SNV tool followed by the ASM tool with “allele-specific methylated site” output format, with an additional SNV quality control filtering step in between them with a custom python script “BS_VCF_filter.py”. ASM output was then used to generate parent-specific CGmaps with another custom python script, “AlleleSpecificCGmapper.py”.

Code used for these methods are available at https://github.com/dawelab/endo-teM-genes

CUT&Tag data have been deposited in the NCBI BioProject database (https://www.ncbi.nlm.nih.gov/bioproject/) under accession number PRJNA874319.

## Acknowledgements

This study was supported in part by resources and technical expertise from the Georgia Advanced Computing Resource Center, a partnership between the University of Georgia’s Office of the Vice President for Research and the Office of the Vice President for Information Technology. This study was funded by NSF grant 2114797 to J.I.G.

## Contributions

Y.S., J.I.G., and Y.Z. analyzed data. D.W.K. and JIG harvested samples and performed CUT&Tag. J.I.G., R.K.D., and Y.S. wrote the paper. J.I.G. and R.K.D. designed and supervised the study.

