## Supplemental Figures for "A switch from TE-like heterochromatin to euchromatin underlies activation of protein storage genes in maize endosperm"

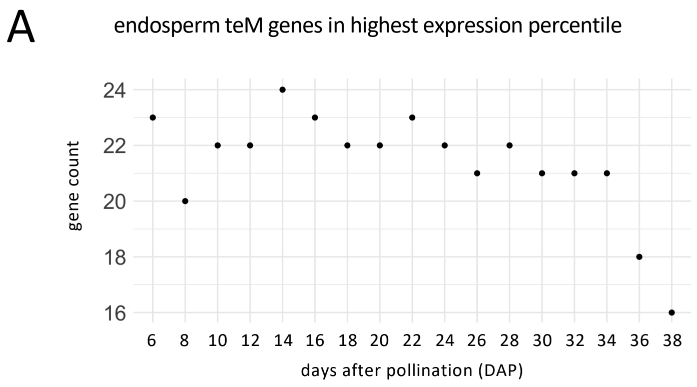


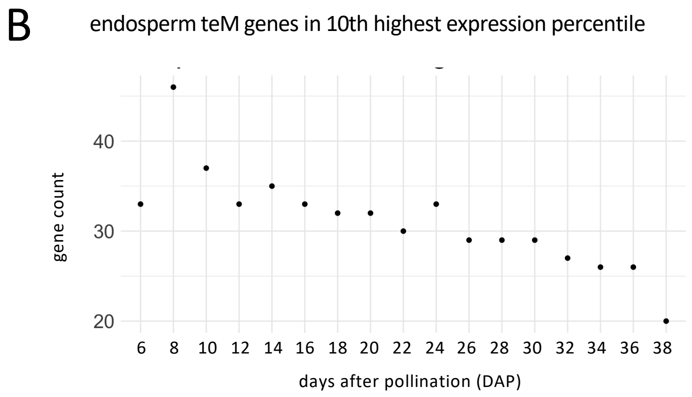


**Supplementary Fig. S1: Endosperm teM genes rank in the top percentiles of gene expression in endosperm.**
The plots show the percent of endosperm teM genes that rank in the top percentile or top tenth percentile of endosperm gene expression at each stage of endosperm development.

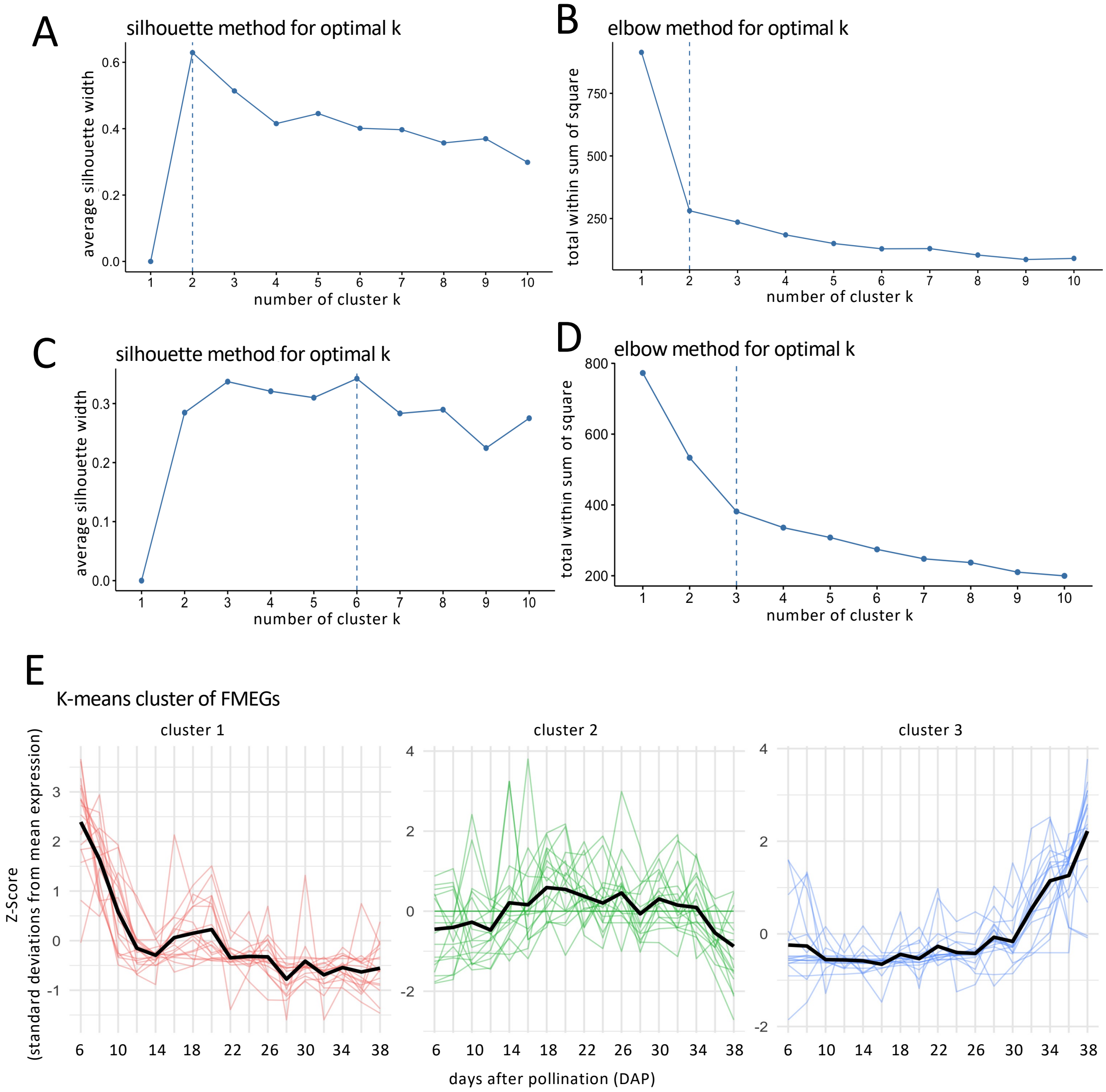


**Supplementary Fig. S2: Clustering of genes by expression over endosperm development**

**a.** Elbow method of determining optimal number of clusters of endosperm teM genes. Vertical dashed line indicates optimal cluster number. **b.** Silhouette method of determining optimal number of clusters of endosperm teM genes. **c.** Elbow method of determining optimal number of clusters of FMEGs. **d.** Silhouette method of determining optimal number of clusters of FMEGs. **e.** Expression patterns based on clustering FMEGs into three groups. The thick black lines indicate the average of all individual genes.

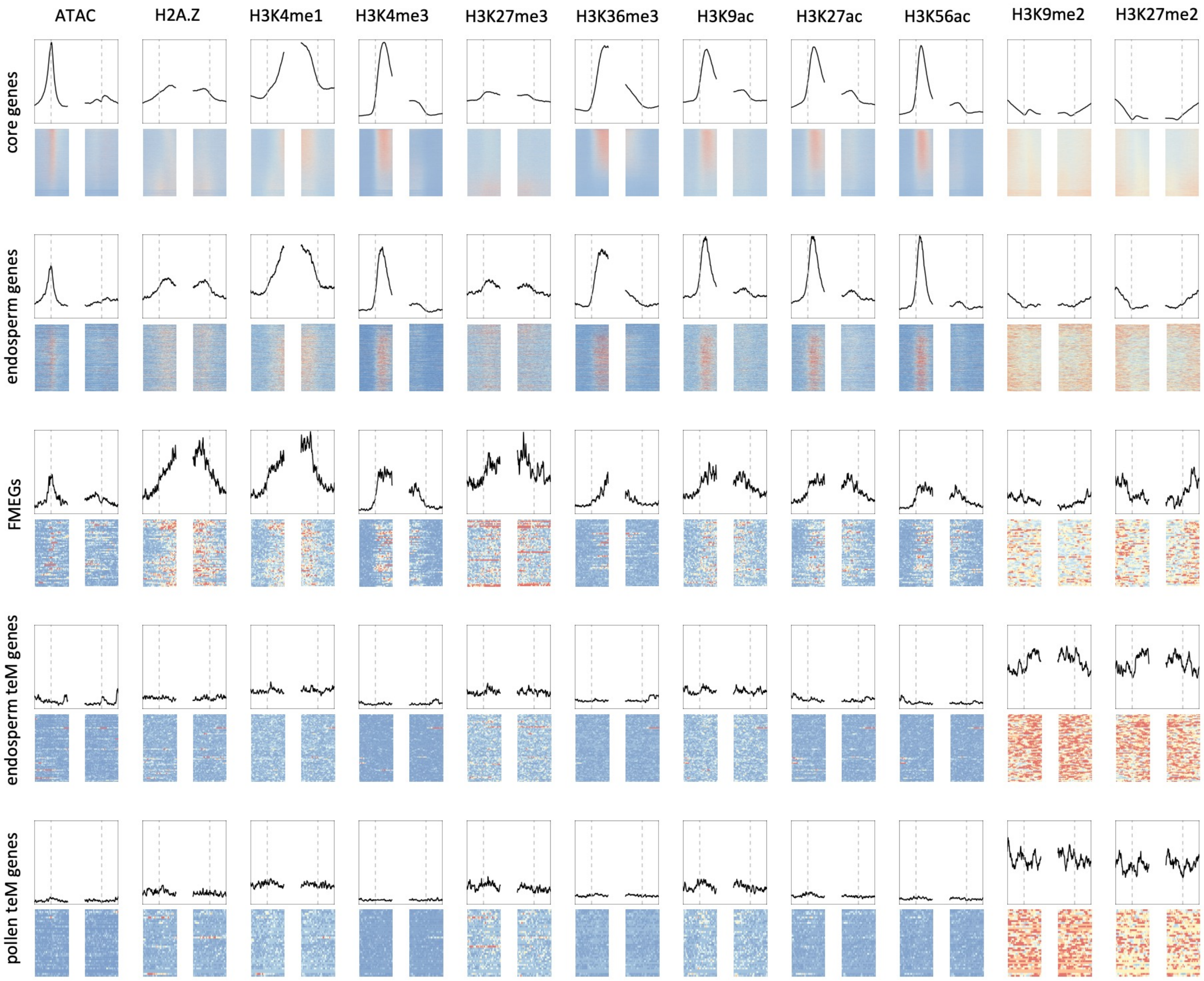


**Supplementary Fig S3. Endosperm and pollen teM genes lack genic chromatin modifications in leaves.**
All heatmaps and metaplots were generated for 2-Kb regions centered on transcription start sites and on polyadenylation sites. Each region was divided into 10-bp bins. For each chromatin feature, the y-axis range of the metagene plots is constant across gene groups and determined by the highest value measured across all gene groups. The color scale of the heatmaps is also constant, with the 95th percentile value for each feature across gene groups used as the maximum color value and zero as the lowest. Genes are ranked by the maximum TPM in endosperm from 6 to 38 DAP [48]. The H2A.Z, H3K27ac, H3K27me3, H3K36me3, H3K4me1, H3K4me3, and H3K56ac, H3K9ac CHIP-seq, and ATAC-seq are from developing second leaves [72]. H3K9me2 and H3K27me2 ChIP-seq was from multiple V4 stage developing leaves [74].

A

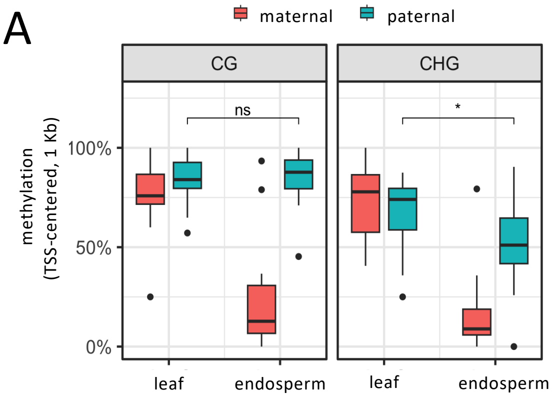


**Supplementary Fig S4. Paternal and maternal CHG methylation is reduced in endosperm.**

Allele-specific methylation was called using genetic variants specific to each genome from EM-seq data from B73 x W22 endosperm and B73 x W22 leaf.

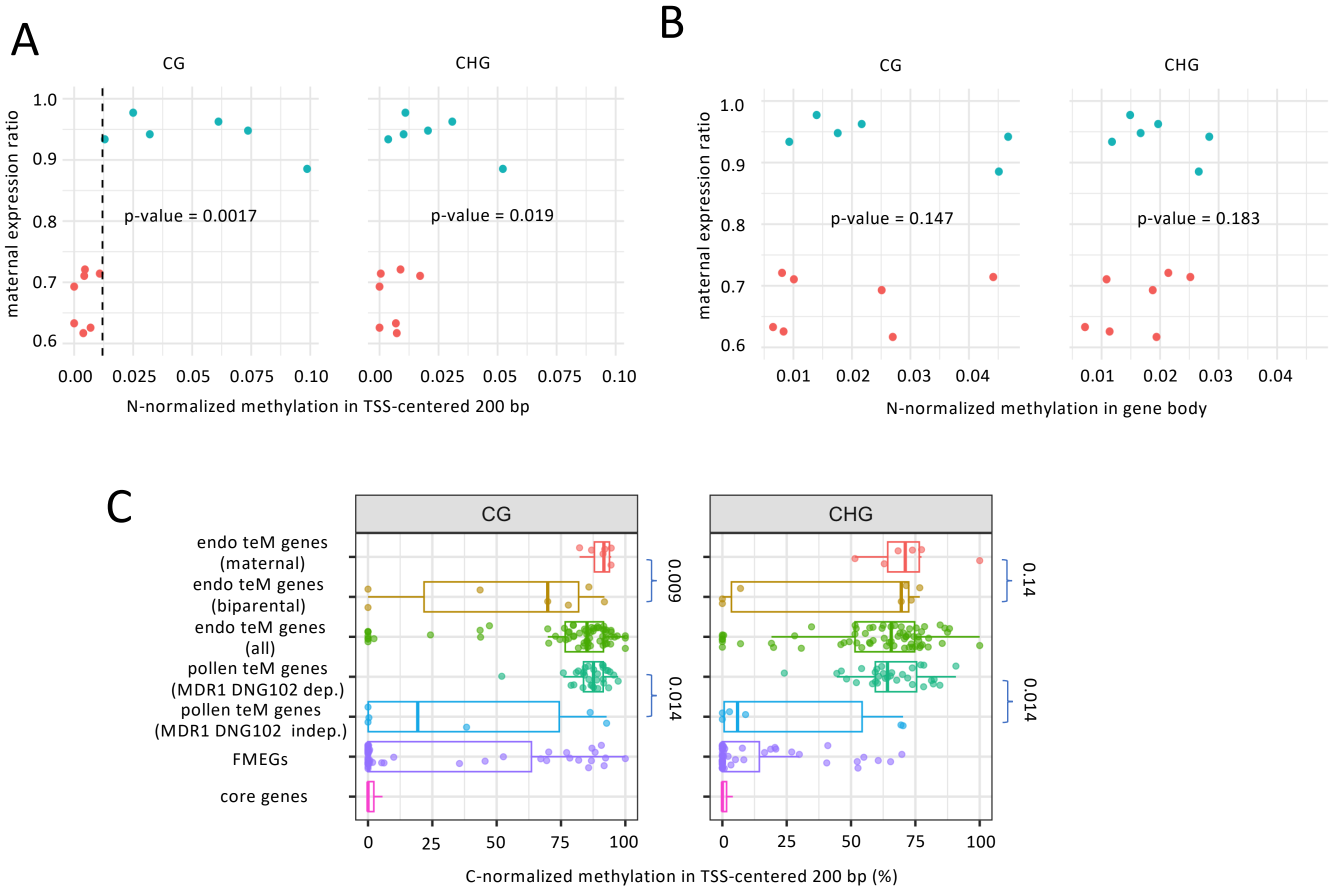


**Supplementary Fig. S5: Methylation of TSSs but not gene bodies correlates with imprinting**
**a.** N-normalized methylation in promoters of endosperm teM genes correlates with their maternal expression ratio. Each dot, colored by imprinting status, corresponds to one of the 13 genes in Fig. 4A. N-normalized methylation is the ratio of methylated cytosines per nucleotide in the 200-bp region, as shown in Fig. 4B. The vertical dashed line indicates the methylation value threshold for predicting maternal preferred expression. P-values are from one-tailed Wilcoxon rank-sum test. **b.** N-normalized methylation in endosperm teM gene bodies correlates with their maternal expression ratio. Each dot corresponds to one of the 13 genes in Fig. 4A. **c.** C-normalized methylation in the 200-bp regions centered on transcription start sites (TSSs) of endosperm teM genes correlates with dependence on demethylation for expression. C-normalized methylation is the ratio of methylated cytosines per total cytosines (in either CG or CHG context) in the 200 bp region. For endosperm teM genes, "maternal" are the six genes in (a) with maternal expression more than threefold greater than expected, "biparental" are the seven with expected maternal expression, and "all" is the complete set of 67. For pollen teM genes, "MDR1 DNG102 dep" are the set of 31 whose pollen expression depends on these two DNGs, and "MDR1 DNG102 indep" are the set of six whose pollen expression is independent of them. Regions with no CGs or CHGs were assigned methylation values of zero rather than excluding them from the analysis. P-values are from one-tailed Wilcoxon rank-sum test.
